# Single-cell phylogenies reveal deviations from clock-like, neutral evolution in cancer and healthy tissues

**DOI:** 10.1101/2022.08.09.503287

**Authors:** Nico Borgsmüller, Monica Valecha, Jack Kuipers, Niko Beerenwinkel, David Posada

## Abstract

How tumors evolve affects cancer progression, therapy response, and relapse. However, whether tumor evolution is driven primarily by selectively advantageous or neutral mutations remains under debate. Resolving this controversy has so far been limited by the use of bulk sequencing data. Here, we leverage the high resolution of single-cell DNA sequencing (scDNA-seq) to test for clock-like, neutral evolution. Under neutrality, different cell lineages evolve at a similar rate, accumulating mutations according to a molecular clock. We developed and benchmarked a test of the somatic clock based on single-cell phylogenies and applied it to 22 scDNA-seq datasets. We rejected the clock in 10/13 cancer and 5/9 healthy datasets. The clock rejection in seven cancer datasets could be related to known driver mutations. Our findings demonstrate the power of scDNA-seq for studying somatic evolution and suggest that some cancer and healthy cell populations are driven by selection while others seem to evolve under neutrality.

## 1 Introduction

Understanding tumor evolution is essential for predicting cancer progression and treatment response [1–5]. Still, the relative role of adaptive and neutral evolution after the malignant transformation has been debated extensively in recent years, without having reached a clear consensus yet [6–11]. Williams *et al.* [7] proposed a model for neutral somatic evolution where multiple clones grow at similar rates after tumor initiation. They used the variant allele frequencies (VAFs) of bulk sequencing samples to test for neutral evolution, assuming that clones with an increased growth rate alter the expected VAF distribution. When applied to samples from different tumor types, they failed to reject neutral evolution in up to 33 % of the datasets analyzed [7, 12]. Several studies, however, questioned these findings and criticized the test proposed by Williams *et al.* as biased, as different simulation approaches or statistical tests lead to contradictory results [9–11].

Alternatively, neutrality can be tested directly by assessing differences in the evolutionary rate among somatic lineages. Under neutral evolution and in a constant environment, different cell lineages accumulate mutations at a constant rate, as in a molecular clock [13,14]. A cell lineage with a growth advantage, in contrast, divides faster and accumulates more mutations per time, leading to distinct evolutionary rates in the cell phylogeny [15]. However, assessing rate heterogeneity among cell lineages with bulk samples is difficult, as millions of cells are sequenced simultaneously. Consequently, cell lineages are mixed, and the deconvolution process is complex and error-prone. Single-cell DNA sequencing (scDNA-seq), in comparison, facilitates the inference of the evolutionary rates among cell lineages [16–19], therefore enabling a direct test of the somatic molecular clock. However, scDNA-seq data suffers from technical errors like false or missed mutations [20] that could bias downstream analysis if not taken into account.

Here, we introduce a Poisson tree (PT) test for detecting deviations from the molecular clock in cell phylogenies inferred from scDNA-seq data. On simulated data, the PT test can identify non-clock evolution while still being robust to scDNA-seq noise. We applied the PT test to 22 scDNA-seq datasets from cancer and healthy tissues and rejected neutral, clock-like evolution in 10/13 cancer and 5/9 normal datasets.

## 2 Results

### 2.1 A Poisson tree (PT) test of the molecular clock

A standard pipeline for scDNA-seq processing includes sampling, isolation, amplification, sequencing, and mutation calling (Fig. 1a). Mutations can be displayed as a mutation matrix (Fig. 1b) and used to infer a cell phylogeny and scDNA-seq error rates (Fig. 1c). The PT test requires as input a mutation matrix, a phylogeny of contemporaneously sampled cells, and error rates. First, it maps mutations to branches and weights them according to their probability of missing true mutations (Fig. 1d). Then, it models the number of mutations per branch with a Poisson distribution and estimates the maximum likelihood under the clock (null hypothesis; Fig. 1e)) and the non-clock (alternative hypothesis; (Fig. 1f)) models. Under the former, the lengths of the branches are constrained, while under the latter, they are independent. Finally, it compares the two models with a likelihood ratio test (LRT) (Fig. 1g). The PT test is described in detail in section 4.1.

**Figure 1:**
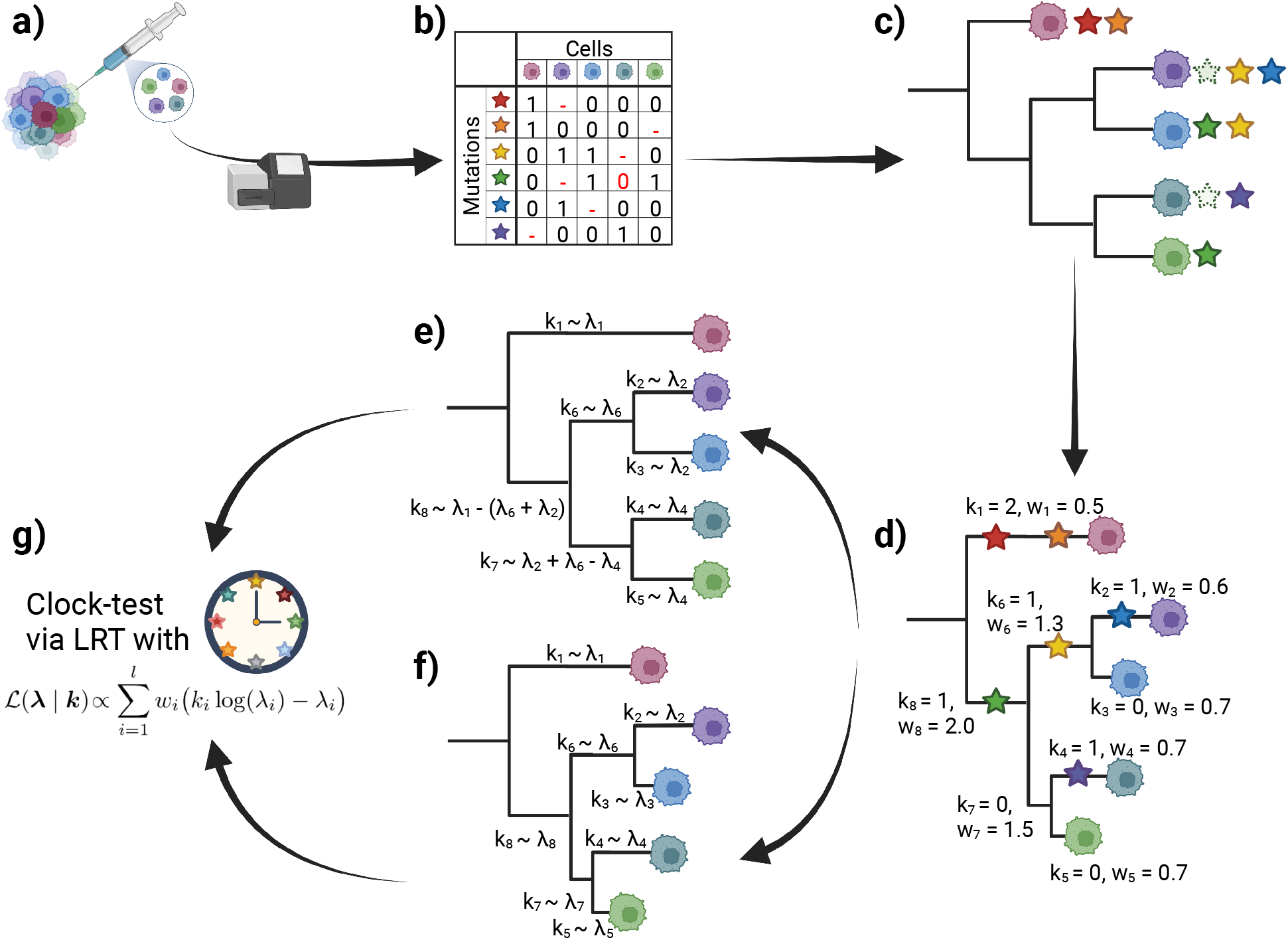
Overview of the Poisson Tree test. Single cells are isolated from a tissue, and their genome is amplified and sequenced (**a**). Based on the sequencing reads, mutations are called and displayed in a mutation matrix (**b**), which is used to infer the cell phylogeny (**c**). Mutations are mapped onto the branches of the cell phylogeny, specifying their length ***k***, and branch weights ***w*** are determined based on the branches’ sensitivity to errors (**d**). Branch lengths ***k*** are modeled by a Poisson distribution with rate parameter **λ**. Under the clock (null) model, the evolutionary rate is constant, implying that the cumulative branch length from the root to any cell is the same (**e**), and the rate parameters **λ** are constrained accordingly. Under the non-clock (alternative) model, the branch lengths are independent and, therefore, unconstrained (**f**). The likelihood of the data under the clock and the unconstrained model is computed and compared with a Likelihood Ratio Test (LRT) (**g**).

### 2.2 The PT test detects clock-deviations reliably

To evaluate the performance of the PT test, we simulated scDNA-seq data under the clock, with and without sequencing errors.

Without scDNA-seq errors, the p-value distribution of the PT test was uniform, as expected for an unbiased test under the null (Fig. 2a, first panel). In the presence of scDNA-seq errors, p-values were strongly shifted towards 1, making the PT test conservative (Fig. 2a, second to fourth panel). With false negative (FN) rates above 0.3, low p-values became more common, indicating that the test may not distinguish between high scDNA-seq error rates and deviations from the clock (Fig. S1). In all cases, the difference in the p-value distributions between using the inferred cell phylogeny and the estimated scDNA-seq error rates using CellPhy [18] (blue) or the true cell phylogeny and simulated scDNA-seq error rates (red) was marginal. Performance was similar when using SCITE [16] instead of CellPhy (Fig. S1).

**Figure 2:**
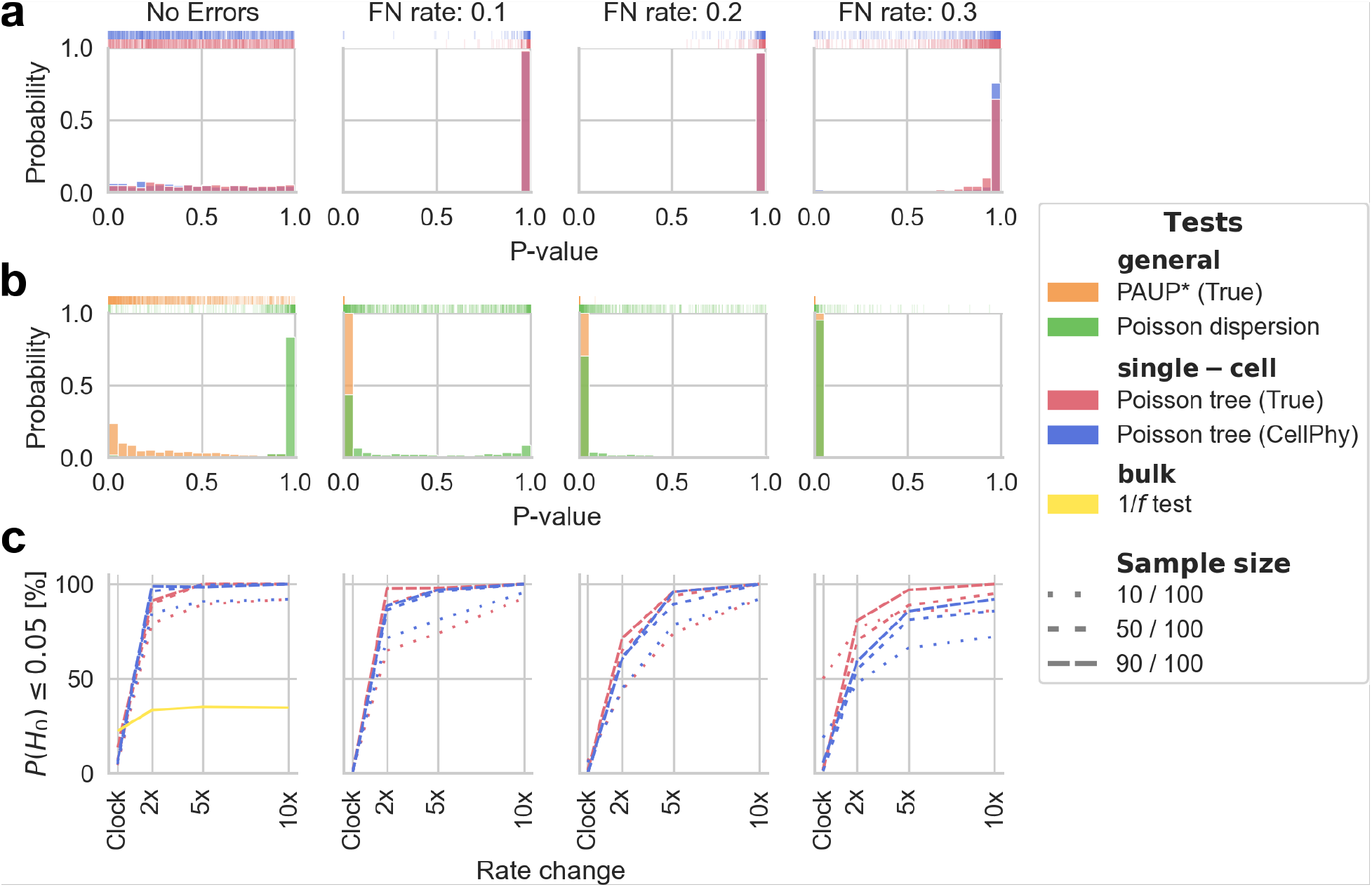
Molecular clock testing on simulated data. **b)** P-value distribution under the clock of the Poisson Tree (PT) test using the true (red) or inferred (blue) cell phylogeny and scDNA-seq error rates, for different scDNA-seq false negative (FN) rates. **a)** P-value distribution under the clock of PAUP*’s LRT (orange) and the Poisson dispersion test (green) for different scDNA-seq FN rates. **c)** Statistical power of the PT test for detecting non-clock evolution. Clock deviations are introduced by changing the evolutionary rate for a given lineage by 2×, 5×, or 10×. Different sample sizes are represented by distinct line styles (total: 100 cells). In the left panel, the yellow line represents the proportion of datasets in which the *1/f* test proposed by Williams *et al.* rejected neutrality on bulk data comparable to the scDNA-seq data.

For comparison, we also applied the clock LRT implemented in PAUP*[21, 22] and the Poisson dispersion test [23]. The former is typically used in organismal phylogenetics and assumes error-free data. The latter tests if the number of mutations per cell is sampled from a Poisson distribution, ignoring the underlying tree topology. The PAUP* LRT was biased towards low p-values, even without scDNA-seq errors and using the true cell phylogeny (Fig. 2b - orange). The p-values of the Poisson dispersion test were biased towards 1 in the absence of scDNA-seq errors (Fig. 2b - green), but became biased towards 0 in the presence of scDNA-seq errors, resulting in high false positive rates. We concluded that both tests are unsuited for testing clock-like evolution with scDNA-seq data.

We also simulated deviations from the clock with CellCoal [24] by introducing changes in the evolutionary rate of a single lineage. For this, we choose a branch with probability proportional to its length. Then, we multiplied its length and that of all descendant branches by 2 ×, 5 ×, or 10 ×. To assess the effect of the sample size, we simulated 100 cells and subsampled 10,30, 50, 70, and 90 cells. As expected, the power of the PT test increased with more drastic evolutionary rate changes and larger sample sizes (Fig. 2c, S3). Without scDNA-seq errors, the power of the PT test was 92 % to 100 % already at the 2× rate changes. With scDNA-seq errors, the power of the PT was above 90 % for 5× and 10× rate changes and samples with more than 10 cells. For the 2× rate change, the power dropped below 50 %, especially for small sample sizes and high error rates. Overall, we conclud that the PT test can reliably assess clock-like evolution in scDNA-seq data.

### 2.3 VAF-based selection tests detect clock-deviations poorly

Next, we simulated bulk data at 100 × depth without scDNA-seq errors, based on the same single-cell phylogenies of the previous section, and ran the 1/*f* test [7] and mobster [25]. We only included datasets where the fraction of cells affected by the rate change was between 20 % and 70 %, as the 1/*f* test detects deviations from neutrality only in that VAF range [25]. The 1/*f* test rejected neutrality in 20 % f the simulations under the clock and in more than 30% of the simulations in the presence of a 2×, 5×, or 10× change of the evolutionary rate (Fig. 2c, first panel - yellow). Mobster did not infer subclonal selection for all evolutionary rate changes (Fig. S2).

### 2.4 The PT test infers clock and non-clock evolution in scDNA-seq data

We applied the PT test to 22 scDNA-seq datasets (10 whole-genome and 12 whole-exome) from 15 patients containing between 7 and 71 cells (Table 1). Thirteen datasets were derived from cancer tissues (blood, bladder, lung, prostate, breast, colorectal (CRC), and renal cancer) and nine from normal, healthy tissue. Additionally, all datasets contained a bulk normal sample and all but two cancer datasets contained a bulk tumor sample. Tables S1 and Spreadsheet S1 describe them in detail. We ran the PT test with the cell phylogenies and scDNA-seq error rates inferred by CellPhy. Then, we mapped the mutations to specific branches (see 4) and identified cancer-specific driver mutations using IntOGen [26].

**Table 1:**
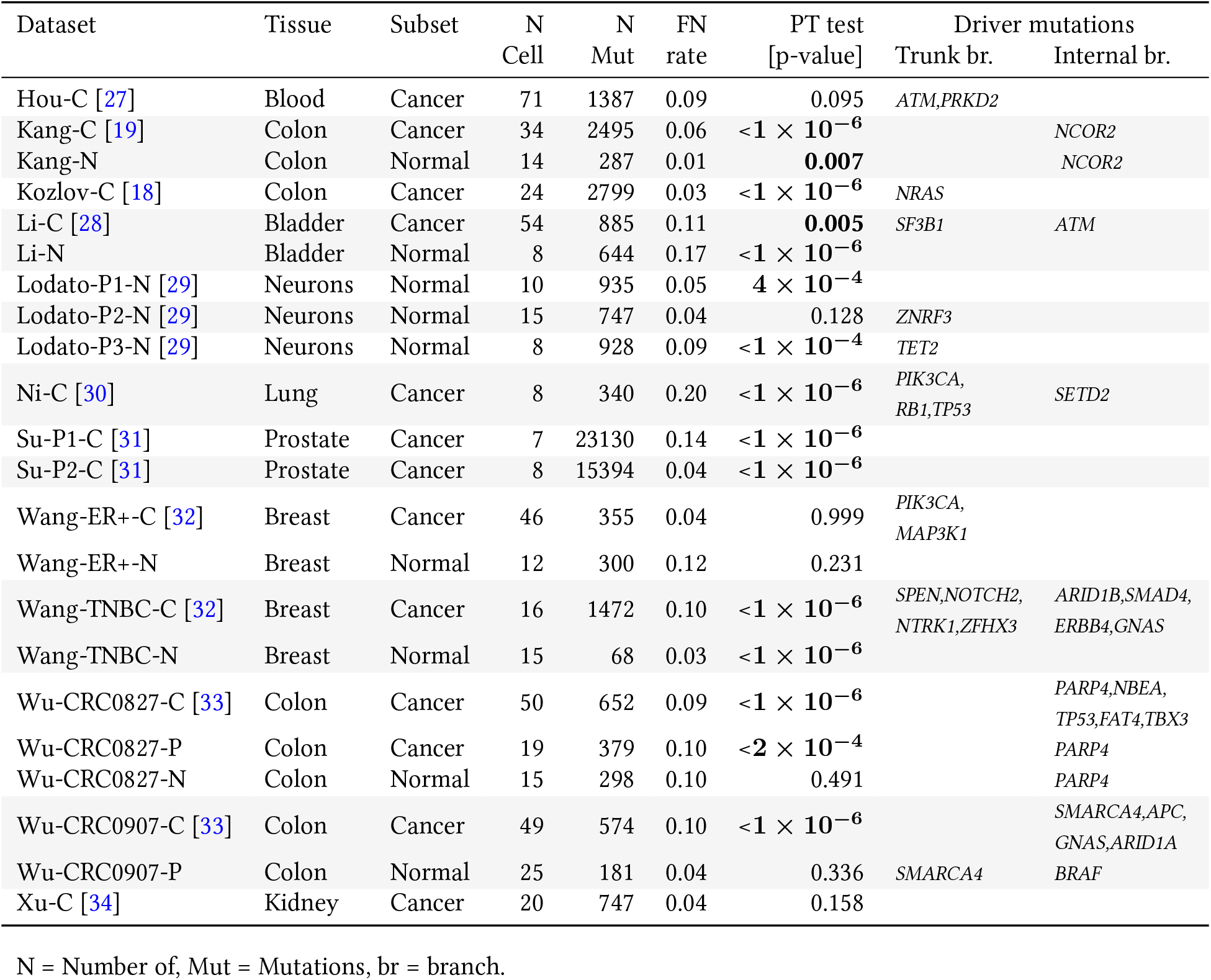
Poisson tree (PT) test results and called cancer-spicific driver genes for scDNA-seq datasets. For the PT test, p-values below a significance level of 0.05 are displayed in bold. N = Number of, Mut = Mutations, br = branch.

Out of the 22 scDNA-seq datasets, we rejected neutral evolution in 10/13 cancer and 5/9 normal data sets. We did not find any relationships between the PT test results and the number of mutations, cells, or inferred scDNA-seq FN rate (Fig. S4). Some of the results of the PT-test might be explained by the identified known cancer-specific driver mutations on internal branches (Fig. 3). Driver mutations on the trunk branch affect all sampled lineages equally and, therefore, will not alter the clock. In four normal datasets (Lodato-P2-N, Wang-ER+-N, Wu-CRC0827-N, and Wu-CRC0907-P), the PT test did not reject the clock (Fig. 3a). In Wang-ER+-N, we detected no driver mutations, and in Lodato-P2-N all known drivers were placed on the trunk branch. In the benign polyp dataset Wu-CRC0907-P, we detected an activating mutation in the oncogene *BRAF* in 3/25 cells, which was not reported in the original study. *BRAF* activation is a known early event in CRC tumor initiation [35], indicating that the polyp might have been adenomatous already. In Wu-CRC0827-N, we inferred a *PARP4* mutation on an internal branch, present in 4/15 cells. However, *PARP4’*s mode of action is labeled as “ambiguous” in IntOGen, and it is not listed as a driver gene in the Cancer Gene Census (CGC) [36]. The PT test rejected the clock in the remaining five normal datasets (Fig. 3b). Within these, only in Kang-N we found a driver mutation on an internal branch (present in 9/14 cells), namely an activation of *NCOR2,* a known driver in the CGC. For Lodato-P1-N, Lodato-P3-N, and Li-N, we inferred fully ladder-like trees, meaning that every internal node was connected to at least one single-cell. Such a pattern might be caused by varying scDNA-seq error rates across otherwise contemporaneous cells. Therefore, these results should be interpreted with caution. In Wang-TNBC-N only 68 mutations were called, out of which 37 were mapped to the trunk branch. The low number of mutations at internal branches did not prevent rejecting of the clock. The PT test did not reject the clock (Fig. 3c) in three cancer datasets (Wang-ER+-C, Hou-C, Xu-C). In all of them, we could not identify drivers on internal branches. For the remaining nine cancer datasets, the PT test rejected the clock (Fig. 3d). In six of these, we identified at least one known driver mutation on an internal branch of the tree.

**Figure 3:**
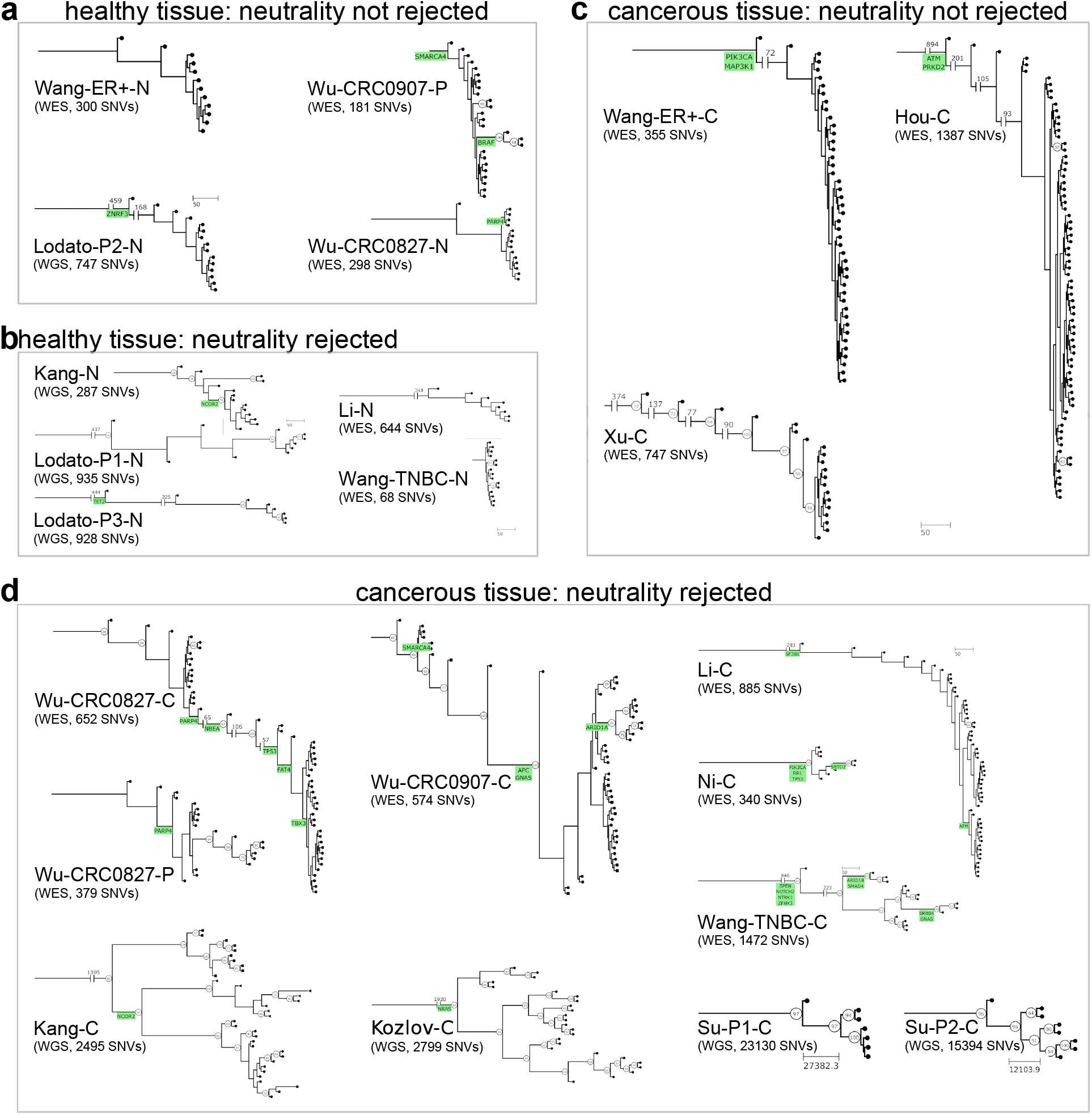
Inferred single-cell phylogenies and known driver mutations. **a)** In four healthy tissue datasets, we detected no deviation from the molecular clock. Driver mutations were either absent, located on the trunk branch, or present in four cells at most. **b)** In five healthy tissue datasets, we rejected the clock. Three of these showed a ladder-like pattern, meaning that each internal node is an ancestor to at least one leaf node (Lodato-P1, Lodato-P3, and Li-N). In the Wang-TNBC-N dataset, we called only 68 mutations and mapped most the trunk branch. In Kang-N, we identified a known driver mutation on an internal branch. **c)** In three cancer datasets, we did not reject the clock. We either identified no driver mutations (Xu-C) or mapped all known drivers to the trunk branch (Hou-C and Wang-ER+-C). **d)** In ten cancer datasets, we found significant deviations from the clock. In seven of these, we identified at least one known driver mutation on an internal branch. In the Kozlov-C datasets, the only known driver was placed on the trunk branch, and in Su-P1-C and Su-P2-C we identified no drivers.

Additionally, we calculated the dN/dS ratios of mutations affecting cancer driver genes for the combined mutations from all individual cells (“pseudo-bulks”) [37]. In ten datasets, mainly derived from normal tissue, the dN/dS ratio could not be calculated as no or just one mutation was located in a cancer driver gene (Table S1). For The leaf node shapes correspond with different spatial sampling locations. Bootstrap values above 50 are indicated. the remaining datasets, the confidence intervals of the dN/dS ratios included 1 (Kang-C, Kozlov-C, Li-C, Li-N, Ni-C, Wang-TNBC-C, Wang-ER+-C, Wang-ER+-N, Wu-CRC0827-C, Wu-CRC0907-C, and Xu-C) or only values smaller than 1 (Hou-C and Su-P1-C). Therefore, the dN/dS ratios showed no evidence for positive selection.

### 2.5 Bulk selection tests produce ambiguous results on scDNA-seq data

Additional to the analysis of the scDNA-seq data, we calculated dN/dS ratios and applied the two approaches by Williams *et al.* to all 16 available bulk tumor samples (Table S2). The dN/dS ratio confidence intervals for all cancer bulk samples included 1. The *1/f* test rejected neutrality in six cases, including two datasets in which the PT test did not (Wu-CRC0907-P and Xu-C). Contrarily, the 1/*f* test did not reject neutrality in four datasets where the PT test did so (Wu-CRC0827-C, Wang-TNBC-C, Ni-C, and Kozlov-C). These findings may be limited as a sequencing depth above 100× and cellularity above 0.5, required for the 1/*f* test to be robust [25], was only achieved in the Li-BC sample. Mobster only produced results for three datasets. In the bladder cancer bulk sample corresponding to Li-C, a clone with selective disadvantage (s = −1.1) was inferred, and no clones were inferred in the bulk samples corresponding to Wang-TNBC-C (PT test p-value: <1e-6) and Xu-C (PT test p-value: 0.16).

## 3 Discussion

The controversies over the mode of tumor evolution arise from the impracticality of directly assessing the effect of individual mutations on cancer progression in humans *in vivo.* Consequently, different tests assess deviations from neutrality indirectly, e.g., via the genome-wide VAF distribution or the ratio of non-synonymous to synonymous mutations in driver genes. In this work, we developed a Poisson tree (PT) test for a molecular clock, implying homogeneity of the evolutionary rates (i.e., the number of mutations accumulated in a time period) among cell lineages. Since we expect clock-like evolution under neutrality [13, 14], deviations from the clock can unveil deviations from neutrality. In somatic evolution, deviations from the clock can result from an increased number of cell divisions per time, i.e., a higher fitness, in a given cell lineage or clone. The expansion of one or more selectively advantageous subclone/s will therefore result in non-clock evolution [15].

Our test is based on single-cell phylogenies, leveraging the high resolution of scDNA-seq data for inferring evolutionary relationships while accounting for the technical noise inherent to scDNA-seq. In our benchmark, the PT test showed a low false positive rate and high power. Minor clock deviations, resulting from effectively neutral evolution, are generally difficult to detect [8], but the PT test was still able to identify variation in the evolutionary rates where the VAF tests did not. When we applied the PT test to 22 real scDNA-seq datasets, we rejected the clock in 15 of them. If the rejection is due to selection, we might be able to locate a driver gene in one of the internal branches of the cell phylogenies. Driver mutations are one of the best understood causes for an increased somatic evolutionary rate [26, 38]. Early driver events, i.e., those mapped to the trunk, will be likely involved in tumor initiation or previous selective sweeps. Later driver events, mapped to internal branches, will result in subclonal selection and a deviation from the clock in the cell phylogeny. In most datasets rejecting the clock, we identified driver mutations on internal branches, possibly causing deviations in the evolutionary rates of different cell lineages. In datasets without clock rejections, we identified either no drivers, drivers on the trunk branch, or drivers being present in only a small fraction of cells. The latter might correspond to late evolutionary rate changes (effectively neutral evolution) but could also result from sampling biases.

We acknowledge that changes in the evolutionary rates, and therefore rejections of the clock, might still occur under neutrality. For example, increases in the mutation rate per cell division for a particular lineage, spatial constraints [39], cell dormancy [40], or variations in the tumor microenvironment of some cells [2], might result in heterogeneous evolutionary rates in the absence of adaptive changes. Indeed, such scenarios might also lead VAF tests (1/*f* test and mobster) to reject “neutrality” [7]. Small sample sizes limit the power of the PT test. Incomplete sampling is an open problem inherent to somatic next-generation sequencing overall, not only to scDNA-seq. In bulk approaches, detectable clones and their VAF distribution depend on the number, type, and spatial location of biopsies taken [41]. While bulk sequencing indeed samples many more cells than single-cell strategies, they rely on summaries of the data that might hide non-obvious levels of evolutionary heterogeneity. For example, although both the PT and VAF tests target recent, ongoing selection, the PT test might be more sensible for newborn selective sweeps.

Understanding whether cell lineages evolve at similar or distinct rates will be relevant for various aspects of cell biology, especially for processes like cancer, development, or differentiation. A molecular clock, for example, is frequently assumed to determine the age of tumors [42], or to study the temporal framework of tissue development [43–45]. Combining established methods from evolutionary biology with advances in single-cell technologies, as we did in this work, offers great potential to study the evolution of somatic tissues through time and space.

## 4 Methods

### 4.1 Poisson tree test model and input data

The Poisson tree (PT) test requires as input a mutation matrix, false positive (FP) and false negative (FN) error rates, and a rooted tree topology 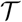 representing the genealogy of the sampled cells. The mutation matrix ***X*** ∈ {0,1, −}^*m×n*^ indicates which of the *m* mutations are present in the *n* cells, with 0 representing the absence of a mutation, 1 the presence, and “–” a missing value. The FP rate *α* is the fraction of 0’s wrongly called as 1’s, while the FN rate *β* is the fraction of true 1’s wrongly called as 0’s. In scDNA-seq data, FPs arise mainly from DNA lesions during cell isolation and manipulations, single-cell whole-genome amplification (scWGA), and sequencing errors; FNs arise mainly from allele dropout (ADO) events during scWGA. Missing values result from ADOs of both alleles or insufficient coverage to call mutations reliably. The fraction of missing data in each cell *j* is 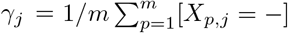, where [·] is the indicator function. The cell tree topology 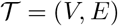 consists of |*V*| = 2*n*–1 nodes and *l* = |*E*| = 2*n* – 2 branches. The n leaf nodes *L* are the sampled cells and the *n* – 1 internal nodes *I* are unobserved ancestor cells. We infer the branch lengths (number of mutations between two nodes) of the cell tree ***k*** = [*k*_1_ … *k_l_*]^*T*^ ∈ ℝ^+^ by mapping mutations to specific branches (see next section). We model the inferred branch length *k_i_* as a Poisson process, depending on a branch-specific evolutionary rate λ_*i*_. In particular, λ_*i*_ ∈ ℝ_>0_ is the product of a mutation rate per cell division and the number of cell divisions.

The likelihood of the evolutionary rates **λ** = (λ_1_,…, λ_*l*_) given the inferred branch lengths ***k*** is then

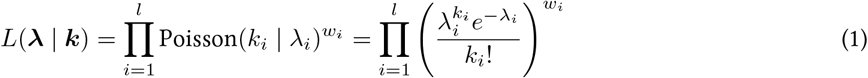

with *w_i_* weighting the impact of branch *i* on the likelihood. If ***w*** = **1** ∈ ℝ^*l*^, all branches are weighted equally; otherwise, branches with a higher weight impact the total likelihood more. The log-likelihood is

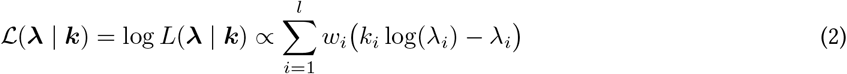

where we have omitted the constant 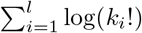, which cancels out in the likelihood ratio below. We infer **λ** by using maximum likelihood estimation (MLE), which amounts to solving

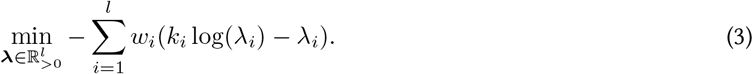

#### Null model H_0_: molecular clock

The null mode assumes a homogeneous evolutionary rate along the cell phylogeny (i.e, a molecular clock) and that all cells have been sampled at the same time point. Consequently, the cumulative branch length from any internal node to its succeeding leaf nodes should be equal. This imposes *n* – 1 constraints on **λ**, which can be written as a system of linear equations defined by a constraint matrix ***C*** ∈ { — 1,0,1}^(*n*-1)×*l*^. Each row in **C** corresponds to an internal node, each column to a branch, and

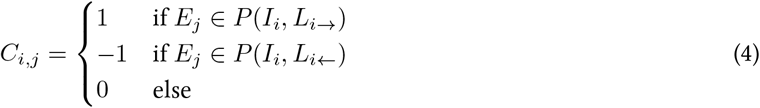

where *P* (*I_x_, L_y_*) is the path between the internal node *x* and the leaf node *y*, i.e., the set of all branches connecting the two nodes, and *L*_*i*→_ and *L*_*i*←_ are arbitrary leaf nodes from the left or right subtree succeeding node *I_i_*, respectively. There are several equivalent parametrizations of the constraint matrix, as both left and right subtrees, as well as the leaf nodes, are chosen arbitrarily. Given that the sum of Poisson-distributed random variables is Poisson-distributed as well, we can write the constraints imposed by the molecular clock as

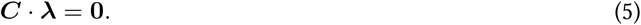

To solve the MLE problem (Eq. 3) subject to the clock constraints (Eq. 5) and the boundary constraints **λ** > **0**, we used the Byrd-Omojokun Trust-Region Sequential Quadratic Programming algorithm [46], a gradient-based numerical optimizer.

#### Alternative model H_1_: constraint-free evolutionary rates

If there are no constraints on **λ**, i.e., **C** = **0**, Eq. 3 can be solved analytically. The likelihood is maximal if the parameters **λ** are equal to the branch lengths ***k*** (Section S2). For branches with no mapped mutations, we use the limit lim_*k*→0+_ *k_i_* log(*k_i_*) = 0.

#### Likelihood ratio test (LRT)

As the null model is nested in the alternative model, their likelihoods can directly be compared with a *χ*^2^-distributed LRT. The test statistic Λ is twice the negative log-likelihood ratio

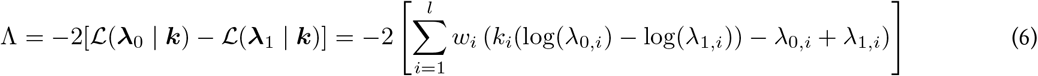

and *χ*^2^-distributed with *n* – 1 degrees of freedom. The probability of the null model given the observed data is

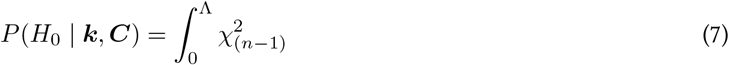

If *k* optimized parameters were on the boundary under the null, however, we changed the distribution of the test statistic to a mixture of *χ*^2^ distributions with *n, n*– 1,…, *n*–*k* degrees of freedom, weighted by normalized binomial coefficients, as reported by Self and Liang [47](*case 9*).

### 4.2 Mapping mutations and defining branch lengths

To map mutations onto specific branches of the cell phylogeny, we define the matrix ***M***^*m*×*l*^ ∈ [0,1], where *M_p,i_* is the probability that mutation *p* is assigned to branch *i*. We assume that mutations are i.i.d. and make the infinite sites assumption (ISA), i.e., we exclude the possibility of parallel and back mutations. Consequently, any mutation that is placed on its true branch is expected to be present in all cells succeeding that branch if no errors occurred. By comparing the expected mutations with the observed ones, we can calculate the probability of assigning a mutation to any branch, similarly to SCITE [16]:

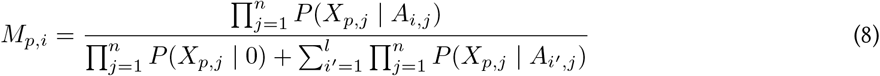

with ***A*** ∈ {0,1}^*l×n*^ being the ancestor matrix representing 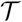, where rows represent branches and columns represent cells, and *A_i,j_* = 1 if branch i belongs to the lineage of cell *j*, and 0 otherwise. The first product of the denominator represents a mutation-free cell and therefore the probability of a wrong mutation call. With FP rate *α* and FN rate *β*, we obtain the following probability for the observed mutation state *x* given an expected state *y*:

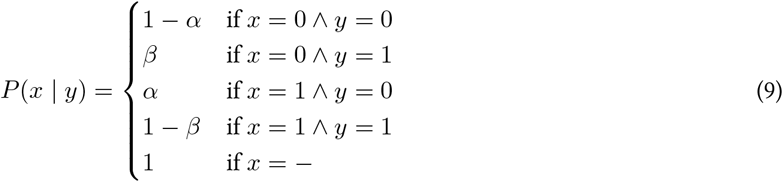

The number of mutations mapped to a branch j is the column sum over **M**:

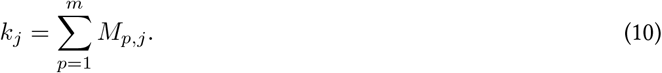

A schematic of the mutation mapping is displayed in Figure S5.

### 4.3 Weighting branches

The estimated branch lengths ***k*** are subject to uncertainties. The soft assignment in Eq. 8 accounts for FP calls and uncertainties in the mutation placement, including FN calls in some but not all cells. However, true mutations can also be not reported in general due to FN or missing events in all cells simultaneously. We call this event a mutation loss. For each branch, the probability of a mutation loss is proportional to the number of cells containing the branch in their lineage, and can be defined as

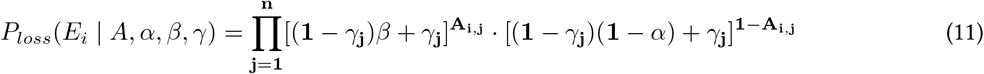

The first term in Eq. 11 describes the probability of FN or missing events in all cells containing the mutation; the second term is the probability of no FP or missing events in any other cell. Given that the probability for a FN call is Bernoulli-distributed, we can weight the branches by their inverse-variance by defining

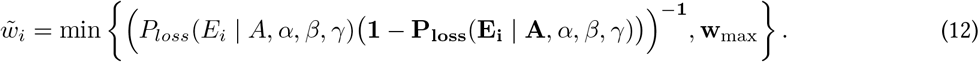

For instance, a *w*_max_ value of 100 corresponds to a maximum probability of 0.99 to observe a true mutation anywhere in the tree, a value of 1000 to a maximum probability of 0.999.

To retain the degrees of freedom of the *χ*^2^ approximation used in the LRT statistic, we normalize the weights

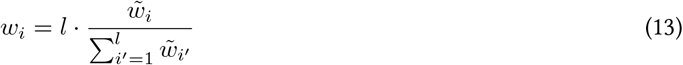

such that 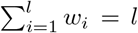. A schematic of the branch weighting is displayed in Figure S6. The limit of the inverse variance of a Bernoulli distribution is infinity, therefore the upper limit *w*_max_ is necessary. Without it, the weight for a single or few branches with a very low probability for mutation losses would be several magnitudes higher than for most other branches. As the weights are normalized, wmax regulates their dispersion: low wmax values lead to weights closer to 1, and larger *w*_max_ values lead to weights dispersed more widely.

We evaluated the impact of *w*_max_ on the PT tests’ accuracy using simulations and found that *w*_max_= 1000 ensured a false positive rate close to zero (Section S1, Fig. S3). Therefore, we used this value for all the calculations in this study.

### 4.4 Simulation of clock and non-clock scDNA-seq data

We used CellCoal [24] to simulate scDNA-seq data with 30 and 100 cells and different levels of error (mainly varying ADO). CellCoal simulates the genealogy of a sample of cells together with genotype data, subject to scDNA-seq errors, in VCF format. To simulate pseudo-bulk data with 100 × depth and a cellularity of 1, we simulated 100 single cells with 1× depth and without scDNA-seq errors. All datasets consisted of 10 000 sites, a somatic mutation rate of 10^-6^, and a sequencing depth of 20 ×. For the simulation of scDNA-seq datasets, the amplification error was 1%, the sequencing error was 1%, and the sequencing depth overdispersion was 5. We increased the ADO rates from 20%± 10% (std) per cell to 80%± 10% per cell in steps of 20 %. As input for the PT test, we used half the ADO rate plus one-third of the amplification error rate as FN. The former represents the chance that the mutated allele is affected by an ADO. Due to the binarization, homozygous mutations are not affected by ADO and heterozygous mutations are affected only in 50 % of the cases. The latter represents the chance of the mutated allele appearing as the reference allele due to a technical error during the scDNA-seq pipeline. As FP rate, we used CellCoal’s amplification error rate. Sequencing errors were ignored, as it is very unlikely that the same sequencing error occurs in multiple reads at the same position. For all datasets, mutation calls with depth < 5 × or quality (GQ) < 1 were filtered out.

By default, CellCoal simulates an ultrametric cell genealogy resulting from a single evolutionary rate (i.e., a clock-like tree). Alternatively, deviations from the clock can be modeled with a single change in the evolutionary rate along the tree. A branch is sampled with probability proportional to its length, and the length of this branch and all descendant branches are multiplied by a given factor.

Here, we simulated non-clock evolution with 2×, 5×, and 10× rate changes. We only included simulations where the fraction of cells affected by the change was between 10 % and 90 %, similar to Williams *et al.*. A smaller fraction corresponds to a late change in the evolutionary rates, a higher fraction to an early change or selective sweep, resulting in effectively neutral, clock-like evolution. To obtain a comparable number of mutations across the different non-clock scenarios, we decreased the global mutation rate for the 2×, 5×, and 10× rate changes to 8^-7^, 5^-7^, and 3^-7^, respectively. Simulations under the null were repeated 1000 times and simulations under the alternative 3000 times.

### 4.5 Inference of cell phylogenies

For the benchmark of the PT test, we inferred the cell phylogenies with CellPhy [18] (default parameters and ML model), which operates a constraint-free model for the branch lengths, and with SCITE [16] (`-r 1 -n 5e5 -d 0.01 -ad 0.2 -e 0.1 -z -a -transpose`), which infers the tree topology but not the branch lengths). Cellphy infers the amplification/sequencing error rate, corresponding to the probability of observing a wrong base, and the ADO rate, rather than FN and FP rates. We used half the ADO rate estimated by CellPhy, plus one-third of the inferred amplification/sequencing error rate, as the FN rate for the PT test. As FP rate, we used the amplification/sequencing error rate inferred by CellPhy. SCITE infers FN and FP rates directly. For the real scDNA-seq data, we inferred cell phylogenies with CellPhy with the same settings as for the benchmark. To root the phylogenetic trees inferred by CellPhy, we added a synthetic cell without any mutation and used it for outgroup rooting.

### 4.6 Statistical tests of neutrality and subclonal selection for bulk data

For the benchmark of the 1/f neutrality test [7], we ran the 1/*f* test (version 0.0.3) with depth 100×, ploidy 2, and cellularity 1. For the benchmark of mobster [48] (version 1.0.0), we ran it with default parameters except for the number of subclones, which we set to 1, according to the single rate change we deployed to simulate deviations from neutrality, resulting in two clades with different evolutionary rates.

For the real bulk data, we used the ploidy and cellularity values inferred by sequenza (version 3.0.0) [49] for the 1/*f* test. Mobster was run with default parameters.

### 4.7 dN/dS ratio estimation

We calculated dN/dS ratios with the R package dndscv [37] and default parameters, including only mutations in the 369 cancer driver genes identified by Martinconera *et al.* [37]. For the scDNA-seq data, we used pseudo-bulk data, as the number of coding mutations per cell was not enough to calculate dN/dS ratios for individual cells.

### 4.8 Biological data processing

All datasets were downloaded in FASTQ format from the NCBI’s Sequence Read Archive (SRA) database. Library adapters and amplification protocol-specific adapters were trimmed with cutadapt (version 1.18). Reads were mapped to the 1000G Reference Genome hs37d5 by using bwa (version 0.7.17), aligned files were sorted by using Picard SortSam (version 2.18.14), and files from different lanes were merged and duplicates marked by using Picard MarkDuplicates (version 2.18.14). GATK IndelRealignement (version 3.7.0) was applied for local realignment based on indel calls by using the 1000G Phase 1 and the Mills and 1000G gold standard databases, and GATK BaseRecalibrator (version 4.0.10) to recalibrate base scores by using dbSNP (build 138) and indels from the 1000G Phase 1. We calculated sequencing depth and breadth with samtools (version 1.9) and the ADO rate as described in [29] for each cell. Cells with extremely high ADO rate (above Q3 + 1.5 IQR per dataset), as well as cells with < 40% coverage breadth, were excluded (Table S2). In contrast to the original studies using WES data, we did not filter the off-target loci but used all sequenced sites. Similar preprocessing was done for the bulk data for normal and tumor samples, followed by estimation of copy numbers using sequenza.

Mutations in single cells were called with a modified version of SCcaller (https://github.com/NBMueller/SCcaller - modifications listed) with default parameters and dbSNP build 138. Additionally, pileups with a minimum mapping quality of 40 were generated with samtools (version 1.9) to call mutations with a modified version of Monovar (https://github.com/NBMueller/MonoVar - modifications listed) with default parameters and without the consensus filtering step. Where tumor bulk samples were available, mutations were called with Mutect2 following the GATK best practice workflow for “Somatic short variant discovery (SNVs + Indels)”. Finally, a set of high-confidence mutations was generated for each dataset by 1) excluding mutations with a quality score below ten or a read depth below ten, and by 2) excluding mutations that were called in only one cell and were not supported by both single-cell callers or by the bulk tumor sample. Additionally, mutations with missing data at more than 50 % of the cells were excluded. To annotate the mutations, we used the Ensembl Variant Effect Predictor [50].

### 4.9 Implementation

The pipelines for processing scDNA-seq data, simulating data, and analyzing data were implemented in Snakemake. The PT test is implemented in Python and requires called mutations in VCF format, a phylogenetic tree in Newick format, and estimated FN and FP rates of the called mutations as input. All the code is freely available at https://github.com/cbg-ethz/scSomMerClock.

## Funding

This work was supported by the European Union’s Horizon 2020 Research and Innovation Program under the Marie Sklodowska-Curie CONTRA grant agreement No. 766030, the European Research Council (ERC) agreements n° 617457 to D.P. and n° 609883 to N.B.], and the Spanish Ministry of Science and Innovation - MICINN (PID2019-106247GB-I00 to D.P.). D.P. also receives support from Xunta de Galicia.

## Conflict of Interest

none declared.

## Notes

https://github.com/cbg-ethz/scSomMerClock

### Competing Interest Statement

The authors have declared no competing interest.

