## Supplementary material for "Single-cell phylogenies reveal deviations from clock-like, neutral evolution in cancer and healthy tissues"

August 9, 2022

### 1 Evaluation of the hyperparameter $w_{\max}$

The power and accuracy of the PT test depend on the hyperparameter $w_{\max}$, defining an upper limit for the branch weights. To assess the effect of $w_{\max}$, we simulated trees with 30 cells and different rate changes and generated ROC curves (Fig. S S3; 3000 repetitions). For all error rates, $w_{\max}$ values $\geq 700$ resulted in false positive rates (FPR) below 0.05, while the true positive rate decreased only marginally for $w_{\max} \geq 700$. To ensure our primary interest, an FPR close to zero, we used a $w_{\max}$ value of 1000. All results on simulated and biological data were all obtained with a $w_{\max}$ value of 1000.

### 2 Maximum likelihood estimate of the alternative model

The likelihood $\sum_{i=1}^{l} w_i \big(k_i \log(\lambda_i) - \lambda_i\big)$ is maximal if $\boldsymbol{\lambda} = \boldsymbol{k}$. This can easily be proven by setting the first derivative equal to zero and solving for $\lambda_i$:

$$\frac{\partial(-\mathcal{L}(\boldsymbol{k} \mid \boldsymbol{\lambda}))}{\partial \lambda_i} = w_i \left(\frac{k_i}{\lambda_i} - 1\right) \overset{!}{=} 0 \quad \Rightarrow \quad \lambda_i = k_i \tag{1}$$

Inserting this solution into the second derivative of the likleihood function verifies that the solution is a maximum:

$$\left.\frac{\partial^2(-\mathcal{L}(\lambda_i \mid \boldsymbol{\lambda}))}{\partial \lambda_i^2}\right|_{\lambda_i = k_i} = -\frac{w_i}{k_i} < 0. \tag{2}$$

If all $\lambda_i$ are i.i.d., then the likelihood is maximal if each $\lambda_i$ is maximal.

| Dataset | Seq. | Ampl. | Tissue | Bulk tumor | Subset | N cell | Avg. depth | Avg. 1× breadth | dN/dS ratio |
|---|---|---|---|---|---|---|---|---|---|
| Hou | WES | MDA | Blood | Yes | Cancer | 71 | 41 | 68 | **0.2 – 0.7** |
| Kang | WGS | Ampli1 | Colon | Yes | Cancer | 34 | 7 | 48 | 0.2 – 6.9 |
| | | | | | Normal | 14 | 8 | 44 | NA |
| Kozlov | WGS | Ampli1 | Colon | Yes | Cancer | 18 | 9 | 50 | 0.1 – 3.0 |
| Li | WES | MDA | Bladder | Yes | Cancer | 54 | 34 | 84 | 0.3 – 2.2 |
| | | | | | Normal | 8 | 35 | 85 | 0.1 – 2.3 |
| Lodato-P1 | WGS | MDA | Neurons | – | Normal | 10 | 55 | 90 | NA |
| Lodato-P2 | WGS | MDA | Neurons | – | Normal | 15 | 43 | 90 | NA |
| Lodato-P3 | WGS | MDA | Neurons | – | Normal | 8 | 51 | 89 | NA |
| Ni | WES | MALBAC | Lung | Yes | Cancer | 8 | 57 | 62 | 0.8 – 4.9 |
| Su-P1 | WGS | MALBAC | Prostate | No | Cancer | 7 | 32 | 54 | **0.0 – 0.5** |
| Su-P2 | WGS | MALBAC | Prostate | No | Cancer | 8 | 28 | 63 | NA |
| Wang-ER+ | WES | MDA | Breast | Yes | Cancer | 16 | 39 | 86 | 0.1 – 4.1 |
| | | | | | Normal | 15 | 37 | 87 | 0.0 – 2.3 |
| Wang-TNBC | WES | MDA | Breast | Yes | Cancer | 46 | 27 | 90 | 0.4 – 1.9 |
| | | | | | Normal | 12 | 25 | 86 | NA |
| Wu-CRC0907 | WES | MDA | Colon | Yes | Cancer | 49 | 55 | 93 | 0.3 – 4.5 |
| | | | | | Polyps | 25 | 58 | 88 | NA |
| Wu-CRC0827 | WES | MDA | Colon | Yes | Cancer | 50 | 69 | 65 | 0.3 – 21.0 |
| | | | | | Polyps | 19 | 61 | 95 | NA |
| | | | | | Normal | 15 | 62 | 94 | NA |
| Xu | WES | MDA | Kidney | Yes | Cancer | 20 | 38 | 75 | 0.2 – 5.1 |

Table 1: Overview of the biological datasets analyzed after filtering. Where the dN/dS ratio is NA, it could not be calculated as no or just one mutation was located in a cancer driver gene. Significance deviations from neutrality are displayed in bold.
Seq. = Sequencing technology, Ampl. = Amplification technology, WES = Whole exome sequencing, WGS = Whole genome sequencing, MDA = Multiple displacement amplification, CRC = Colorectal cancer.

| Dataset | Subset | Depth | Cellularity | *1/f* test | Mobster $s$ | dN/dS ratio |
|---|---|---|---|---|---|---|
| Kang-BC | Primary | 46.5 | 0.99 | **0.001** | no tail | 0.67 – 4.3 |
| Kozlov-BC1 | Tumor Inferior | 42.9 | 0.92 | 0.058 | no tail | 0.77 – 3.1 |
| Kozlov-BC2 | Tumor Middle | 39.8 | 0.87 | 0.92 | no tail | 0.77 – 3.1 |
| Hou-BC1 | LC | 54.2 | 1 | 0.941 | no tail | 0.37 – 2.7 |
| Hou-BC2 | LN | 35.9 | 0.99 | 0.785 | no tail | 0.37 – 2.7 |
| Li-BC | Primary | 176.1 | 0.91 | 0.529 | **−1.096663** | 0.24 – 3.8 |
| Ni-BM | Metastasis | 51 | 0.91 | 0.184 | no tail | 0.63 – 5.9 |
| Ni-BC | Primary | 35.5 | 0.19 | 0.282 | no tail | 0.63 – 5.9 |
| Wang-TNBC-BC | Primary | 71.6 | 0.34 | 0.215 | no tail | 0.74 – 2.4 |
| Wang-ER+-BC | Primary | 86.5 | 0.97 | **0.03** | 0 | NA |
| Wu-CRC0907-BC | Cancer | 51.4 | 0.33 | **0.00** | no tail | 0.22 – 2.6 |
| Wu-CRC0907-BP | Polyps | 48.3 | 0.28 | **0.02** | no tail | 0.22 – 2.6 |
| Wu-CRC0827-BC1 | C | 25 | 0.1 | 0.061 | no tail | 0.05 – 1.7 |
| Wu-CRC0827-BC2 | CA | 29.9 | 0.1 | 0.422 | no tail | 0.05 – 1.7 |
| Wu-CRC0827-BP | Polyps | 58.8 | 0.44 | **0.001** | no tail | 0.05 – 1.7 |
| Xu-BC | Primary | 126.8 | 0.4 | **0.001** | 0 | 0.09 – 11 |

Table 2: Bulk test results for scDNA-seq datasets. Significance deviations from neutrality are displayed in bold.

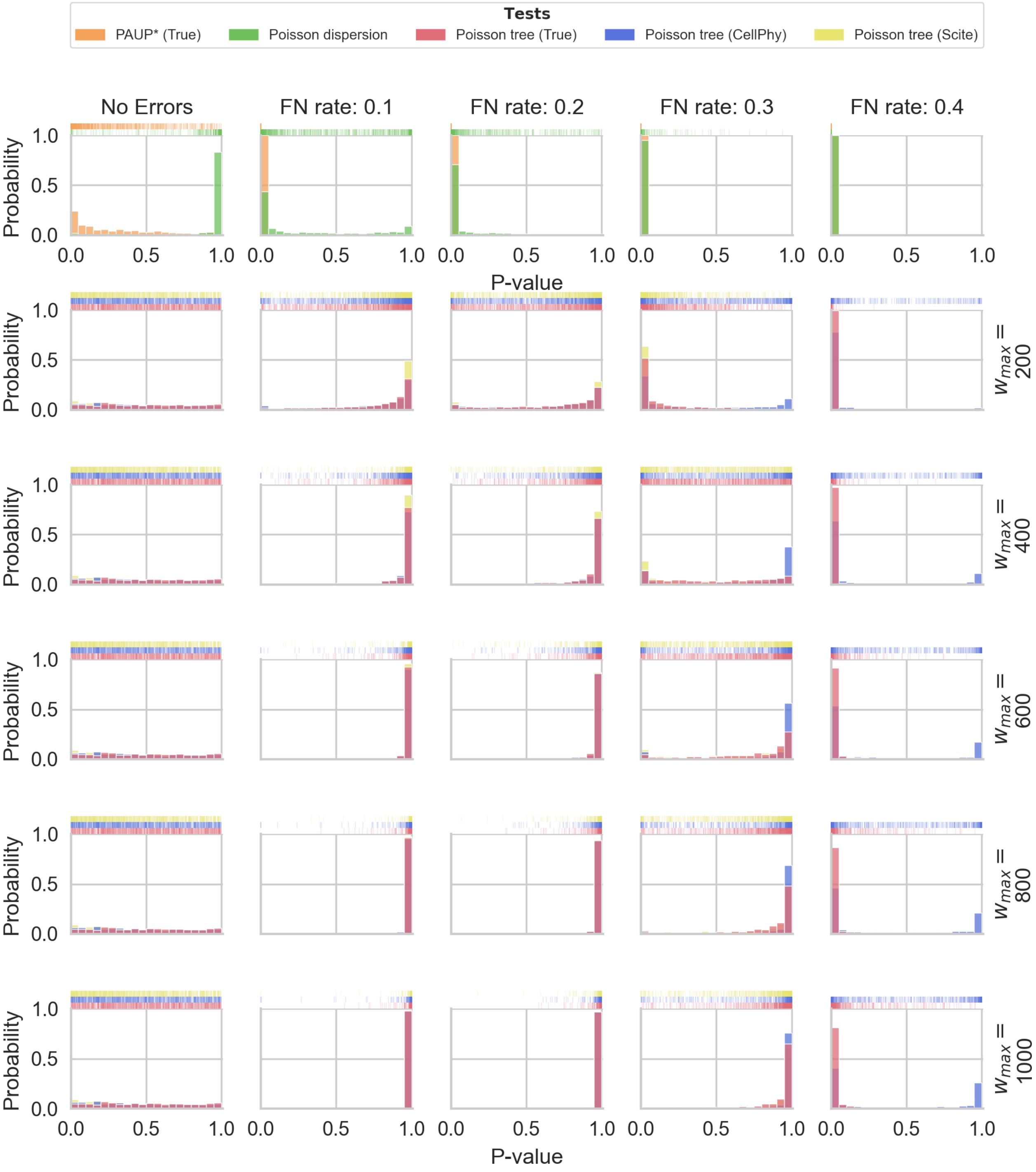


Figure S1: P-value distribution under the clock, for different scDNA-seq false negative (FN) rates (columns) and $w_{\max}$ values (rows). Without errors, the p-values are uniformly distributed. With errors, p-values are shifted towards 1. The higher the $w_{\max}$ value, the stronger the shift. Increasing error rates enhance low p-values (again).

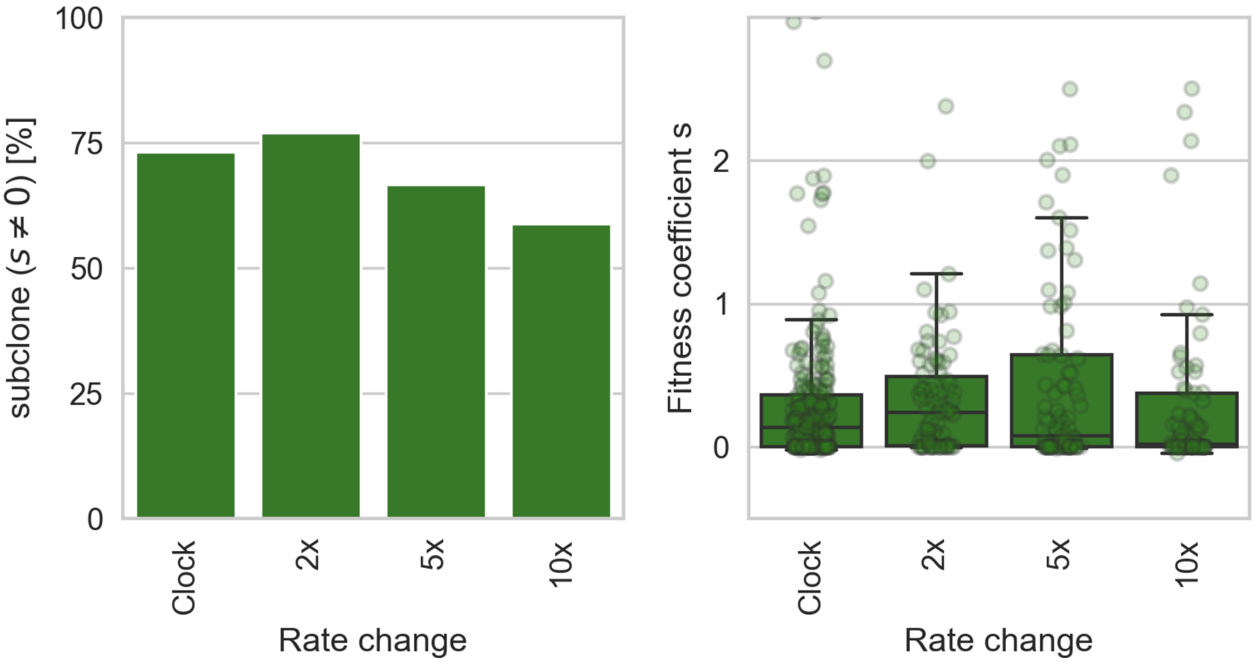


Figure S2: Performance of mobster on bulk simulations (500 repititions). The first panel shows how often mobster inferred a subclone with selective advantage. Under the null, it wrongly inferred a selectively advantageous subclone in 75 % of the cases. In the presence of a lineage with increased evolutionary rate, this value increased only slightly for small rate changes (2×) and dropped to 60 % for larger rate changes. The second panel shows the inferred fitness coefficient of subclones with selective advantage. Under the null, the inferred fitness is small (Median: $< 0.2$). Under the alternative, the fitness coefficients increase slightly for 2× but decreases for higher rate changes. However, more extreme values are fitness coefficients are inferred for higher rate changes (note that the y-axis is cut and does not display all data points).

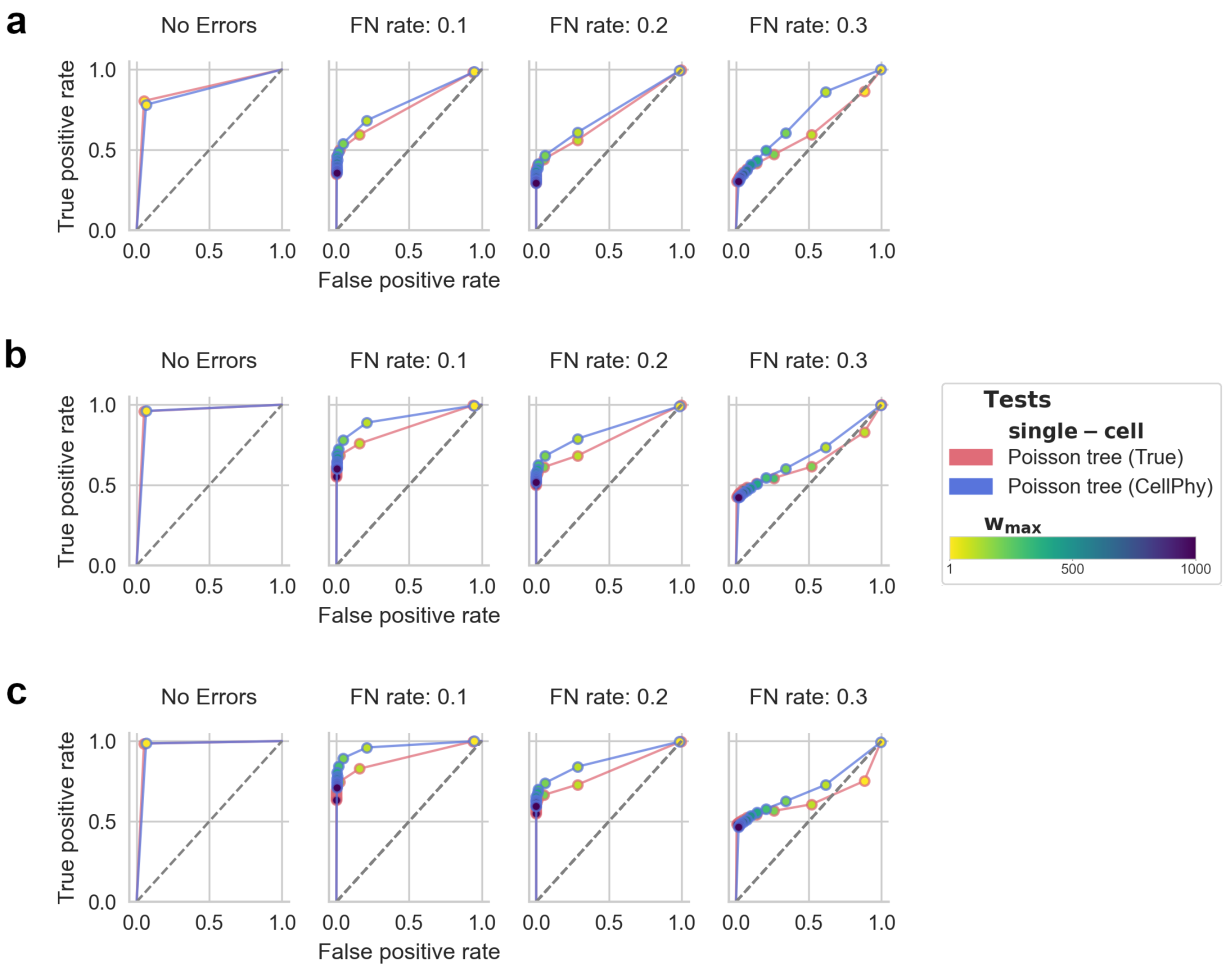


Figure S3: ROC curves of the PT test in dependence of the hyperparameter $w_{\max}$, defining the upper limit of the branch weights. The evolutionary rate change is **a)** 2×, **b)** 5×, and **c)** 10×. With higher $w_{\max}$ values, the PT test became more conservative, and true positive and false positive rates decreased, accordingly. For a clock test, we are interested in low false positive rates, thus, favoring high $w_{\max}$ values.
30 cells simulated, 3000 repetitions, and significance level $\alpha = 0.05$.

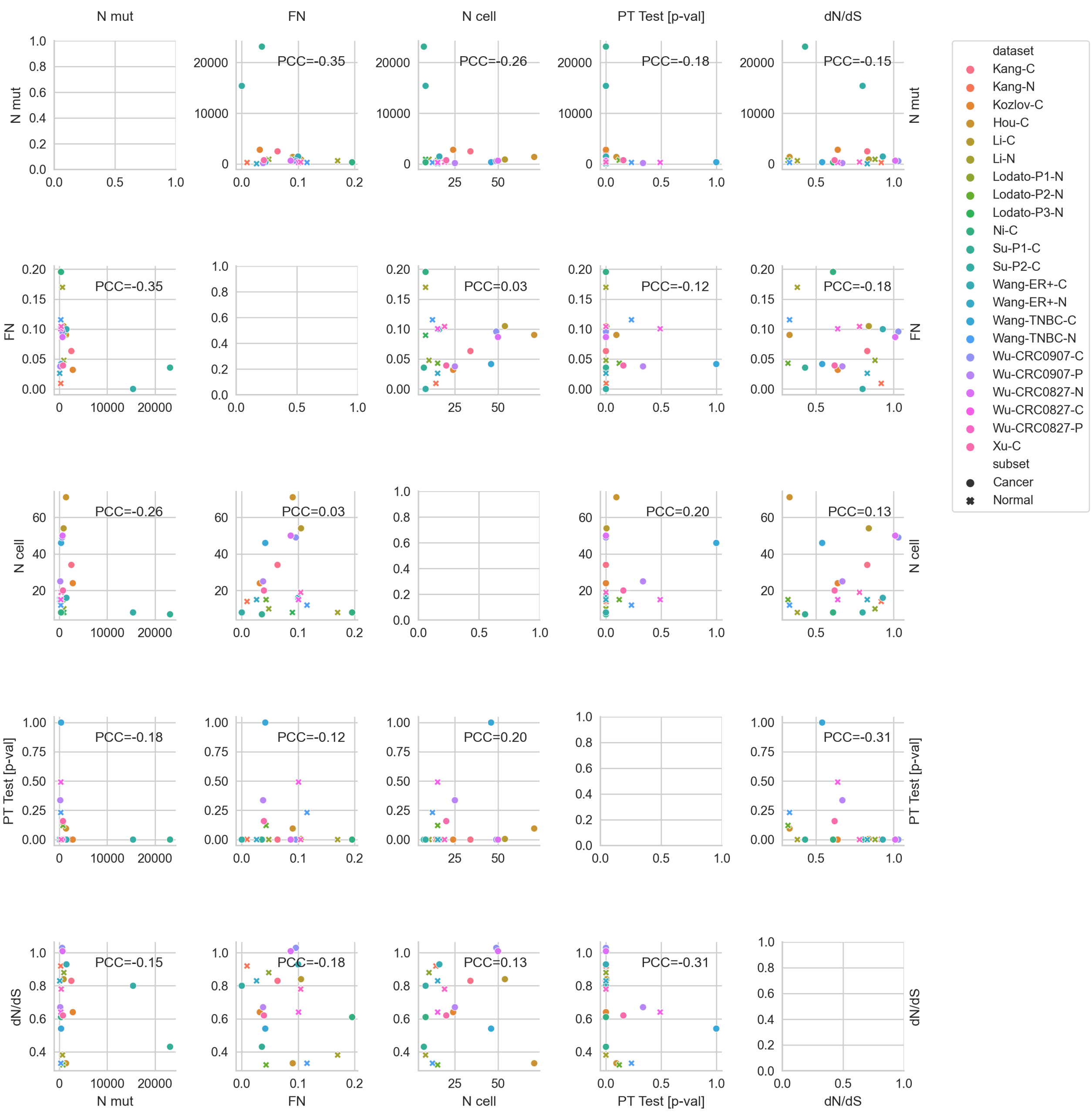


Figure S4: Correlations between features in the biological data. PCC = Pearson Correlation Coefficient.

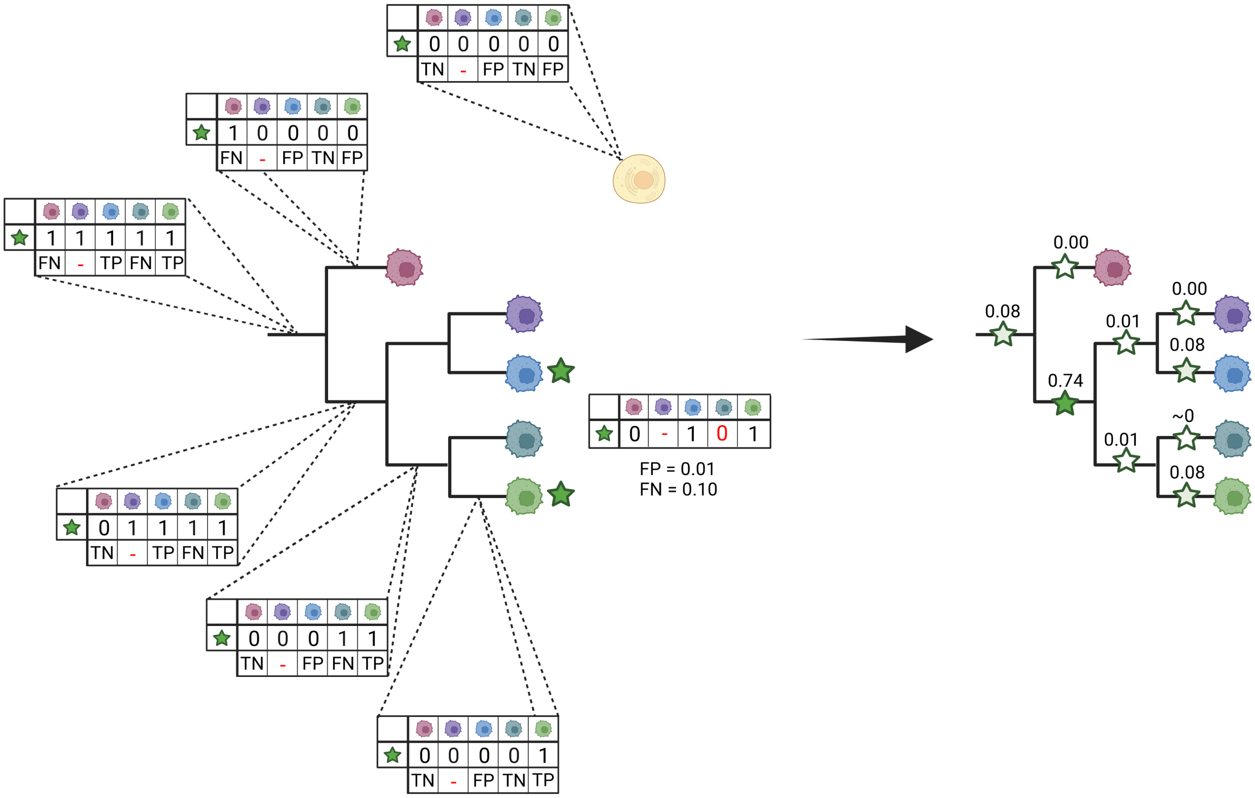


Figure S5: Schematic of the mutation mapping.

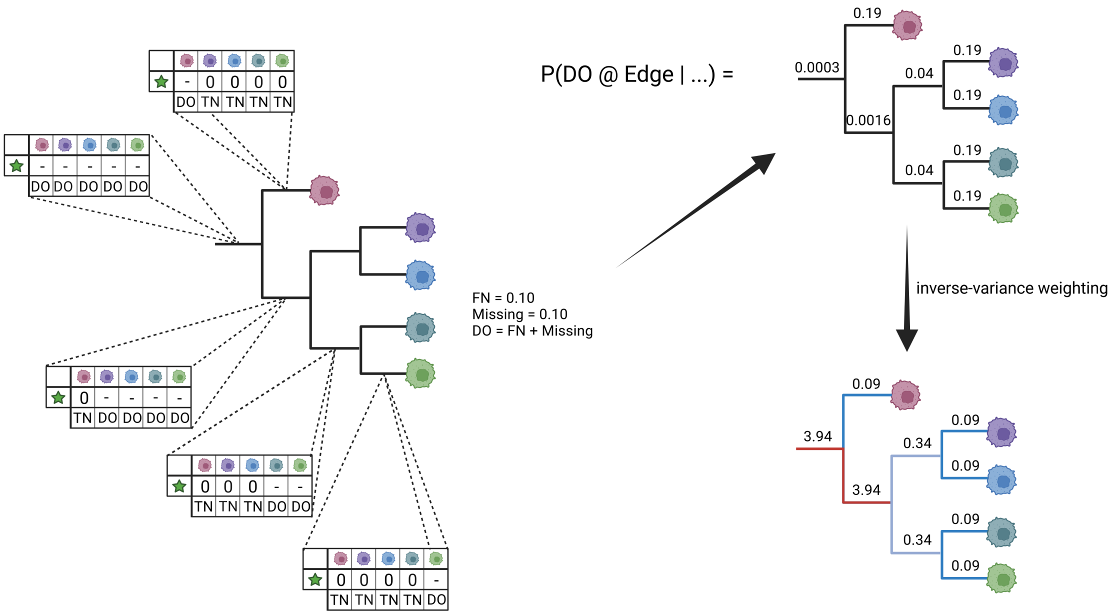


Figure S6: Schematic of the branch weighting.
